# IFNγ activates an immune-like regulatory network in the cardiac vascular endothelium

**DOI:** 10.1101/2024.05.03.592380

**Authors:** Timothy D. Arthur, Isaac N. Joshua, Jennifer P. Nguyen, Agnieszka D’Antonio-Chronowska, Matteo D’Antonio, Kelly A. Frazer

**Affiliations:** Biomedical Sciences Graduate Program, University of California, San Diego, La Jolla, 92023; Department of Biomedical Informatics, University of California, San Diego, La Jolla, 92023; Institute of Genomic Medicine, University of California, San Diego, La Jolla, 92023; Bioinformatics and Systems Biology Graduate Program, University of California, San Diego, La Jolla, 92023; Center for Epigenomics, University of California, San Diego, La Jolla, 92023; Department of Pediatrics, University of California, San Diego, La Jolla, 92023

## Abstract

The regulatory mechanisms underlying the response to pro-inflammatory cytokines in cardiac diseases are poorly understood. Here, we use iPSC-derived cardiovascular progenitor cells (CVPCs) to model the response to interferon gamma (IFNγ) in human cardiac tissue. We generate RNA-seq and ATAC-seq for four CVPCs that were treated with IFNγ and compare them with paired untreated controls. Transcriptional differences after treatment show that IFNγ initiates an innate immune cell-like response, shifts the CVPC transcriptome towards coronary artery and aorta profiles, and stimulates expression of endothelial cell-specific genes. Analysis of the accessible chromatin shows that IFNγ is a potent chromatin remodeler and establishes an IRF-STAT immune-cell like regulatory network. Finally, we show that 11 GWAS risk variants for 8 common cardiac diseases overlap IFNγ-upregulated ATAC-seq peaks. Our findings reveal insights into IFNγ-induced activation of an immune-like regulatory network in the cardiac vascular endothelium and the potential role that regulatory elements in this pathway play in common cardiac diseases.

**Graphical Abstract:** 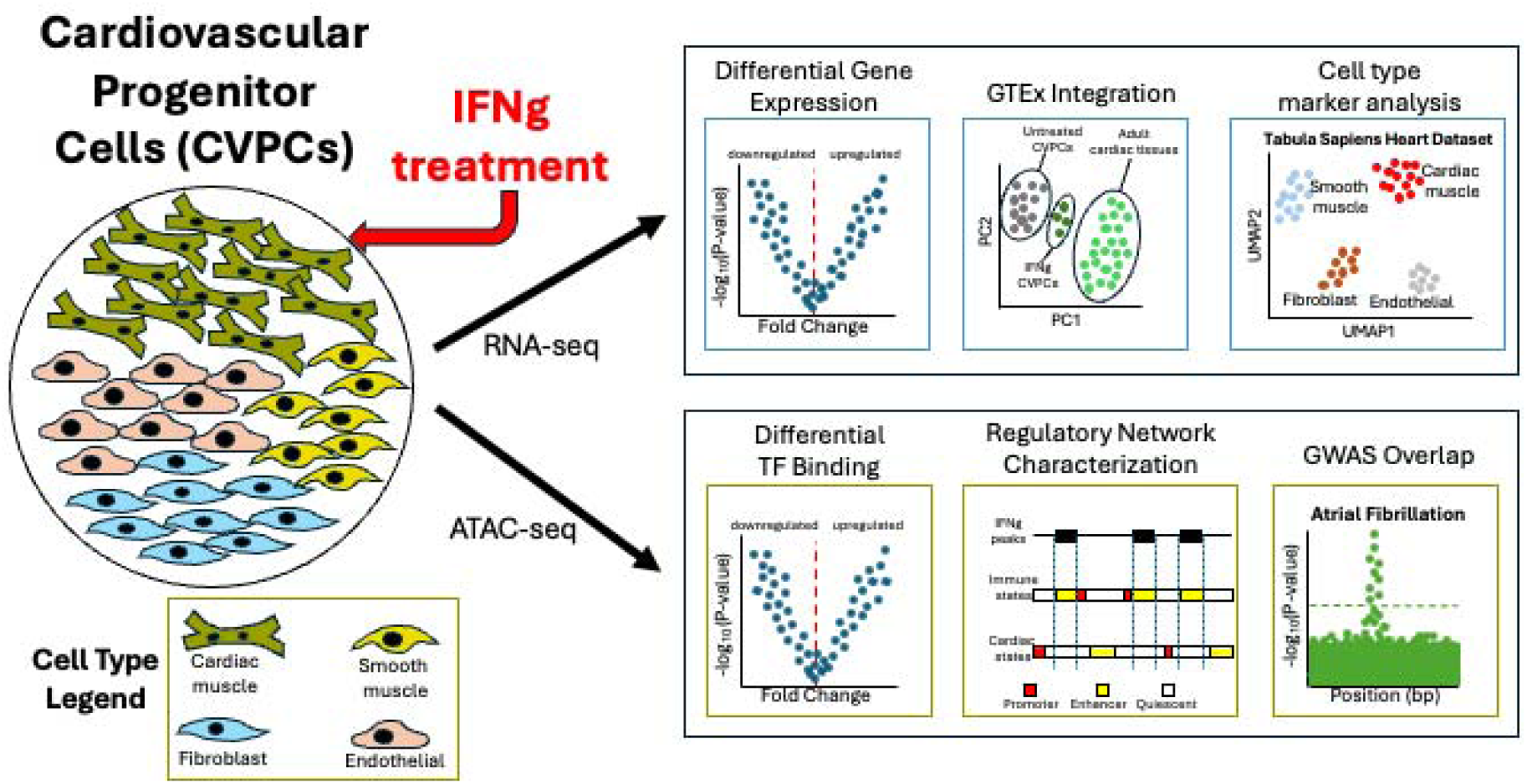

Paired RNA-seq and ATAC-seq was generated for induced pluripotent stem cell derived cardiovascular progenitor cells (CVPCs) treated with interferon-gamma (IFNγ) and matched controls to model the effect of the pro-inflammatory cytokine on human cardiac tissue. Using the RNA-seq, transcriptomic changes were characterized by performing differential gene expression analysis, integrating gene expression data from hundreds of samples of adult cardiac tissues, and using single cell RNA-seq to evaluate cell type-specificity. Using the ATAC-seq, epigenomic changes were characterized by performing differential chromatin accessibility and transcription factor binding analyses, and annotating ATAC-seq peaks with chromatin states from over 800 tissues. Genetic variants in risk loci associated with cardiac diseases were intersected with ATAC-seq peaks to evaluate whether they were in chromatin that is only accessible after IFNγ treatment. The findings demonstrate the utility of using CVPCs to model the effects of cytokines on cardiac tissues and provide the framework for conducting large scale studies to further evaluate GWAS loci that are explained by context-specific regulatory variation.

## Introduction

Cardiovascular disease (CVD) remains a leading cause of morbidity and mortality worldwide, encompassing a broad spectrum of conditions that lead to heart failure, arrhythmias, and vascular dysfunction. Inflammatory cytokines play a central role in the pathophysiology of many cardiac diseases [1–3], with interferon gamma (IFNγ) emerging as a key modulator of the inflammatory response. IFNγ is primarily produced by activated T cells and natural killer (NK) cells and mounts protective responses against pathogens but also has context-specific effects on different cardiac cell types. For example, IFNγ activates the immune JAK-STAT pathway in the vascular endothelial cells in the heart during viral infection[4,5], but damages cardiomyocyte mitochondria by promoting oxidative and nitrosative stress [6]. Increased levels of IFNγ have been observed in ischemic heart disease [7,8], dilated cardiomyopathy [3], and has been identified as a reliable predictor of strokes in patients with atrial fibrillation [1,9]. Given its diverse roles and implications in a myriad of cardiac conditions, comprehending the specific cellular effects of IFNγ remains a significant challenge.

Most efforts to understand the role of IFNγ in cardiac disease pathologies have relied on mouse models [10–12] because obtaining primary cardiac tissues from patients is difficult and often not possible. Induced pluripotent stem cell (iPSC) derived cardiovascular progenitor cells (CVPCs) are a promising model that enables the evaluation of cardiac tissue’s response to stimuli and cardiac disease mechanisms because they can be derived directly from patients without obtaining a direct biopsy from the heart. The iPSCORE collection consists of 180 CVPCs that were derived from iPSCs from 139 individuals using a lactate selection protocol that only gives rise to cardiac cell types [13]. These CVPCs have extensive molecular characterization, including RNA-seq [14,15], ATAC-seq [16], H3K27 acetylation ChIP-seq [16], ChIP-seq for NKX2-5[17], and Hi-C [18]. In previous studies, we have demonstrated that CVPCs exhibit a fetal-like transcriptomic profile and are composed of several cardiac cell types, including cardiac muscle, smooth muscle, endothelial, and fibroblast cells [14], which enables the characterization of cell type-specific drug responses and contributions to disease. Previous studies have shown that, despite the phenotypic differences between adult and fetal cardiac tissue, they both respond similarly to IFNγ [2,19,20], suggesting that fetal-like CVPCs are suitable model for investigating cell-type specific responses.

Here, we use CVPCs from the iPSCORE collection to model the response to IFNγ in cardiac tissue. We generate RNA-seq and ATAC-seq for four CVPCs derived from iPSCs from four unrelated individuals that were treated with IFNγ and compare them with paired untreated controls. We show that in all four CVPC lines, IFNγ treatment shifts the CVPC transcriptome towards coronary artery and aorta-like tissues from the Genotype-Tissue Expression (GTEx) Consortium [21]. Integration of single-cell RNA-seq generated from human heart tissue [22] reveals that IFNγ treatment primarily upregulates endothelial-specific genes. We also show that IFNγ is a potent chromatin remodeler, and binding sites for IRF and STAT family transcription factors are significantly enriched in the differentially accessible chromatin. We next demonstrate that IFNγ activates an innate immune cell-like regulatory network in vascular endothelial cells. Finally, we show that 8 IFNγ-upregulated ATAC-seq peaks overlap 11 GWAS variants for 8 cardiac diseases, suggesting that certain GWAS loci may capture context-specific regulatory variation that are only active after IFNγ stimulation.

Together, this study highlights the utility of CVPCs for modeling the transcriptomic and epigenomic effects of cytokine stimulation and provides key insights into cardiac disease mechanisms. Additionally, this approach can inform *in vivo* experiments and serve as a framework for larger scale studies that can evaluate cell type-specific and genetic effects at higher resolutions.

## Results

### IFNγ stimulates immune cell-independent inflammatory response in cardiac tissue

To examine the effects of IFNγ on gene expression in cardiac tissues, we performed RNA-seq on four CVPCs that were stimulated with IFNγ and four paired untreated controls (Table S1). We performed differential expression and identified 2,729 differentially expressed genes (FDR < 0.05), including 1,594 significantly upregulated by IFNγ stimulation and 1,135 that were downregulated (Figure 1a). Despite the absence of immune cell types in CVPCs, many of the upregulated genes were members of the pathogen-responsive JAK-STAT pathway [23] such as *STAT1* (Figure 1b), *IRF1*, *IFI35*, and *ISG15* (Figure 1a, Table S2). Concordant with previous findings showing that inflammation increases PD-L1 levels in endothelial cells [12], we observed upregulation of *CD274*, which encodes for PD-L1 (Figure 1b). Interestingly, *LEFTY2* and *DANCR*, genes involved in early developmental processes [24,25], are among the genes most downregulated (Figure 1b).

**Figure 1.**
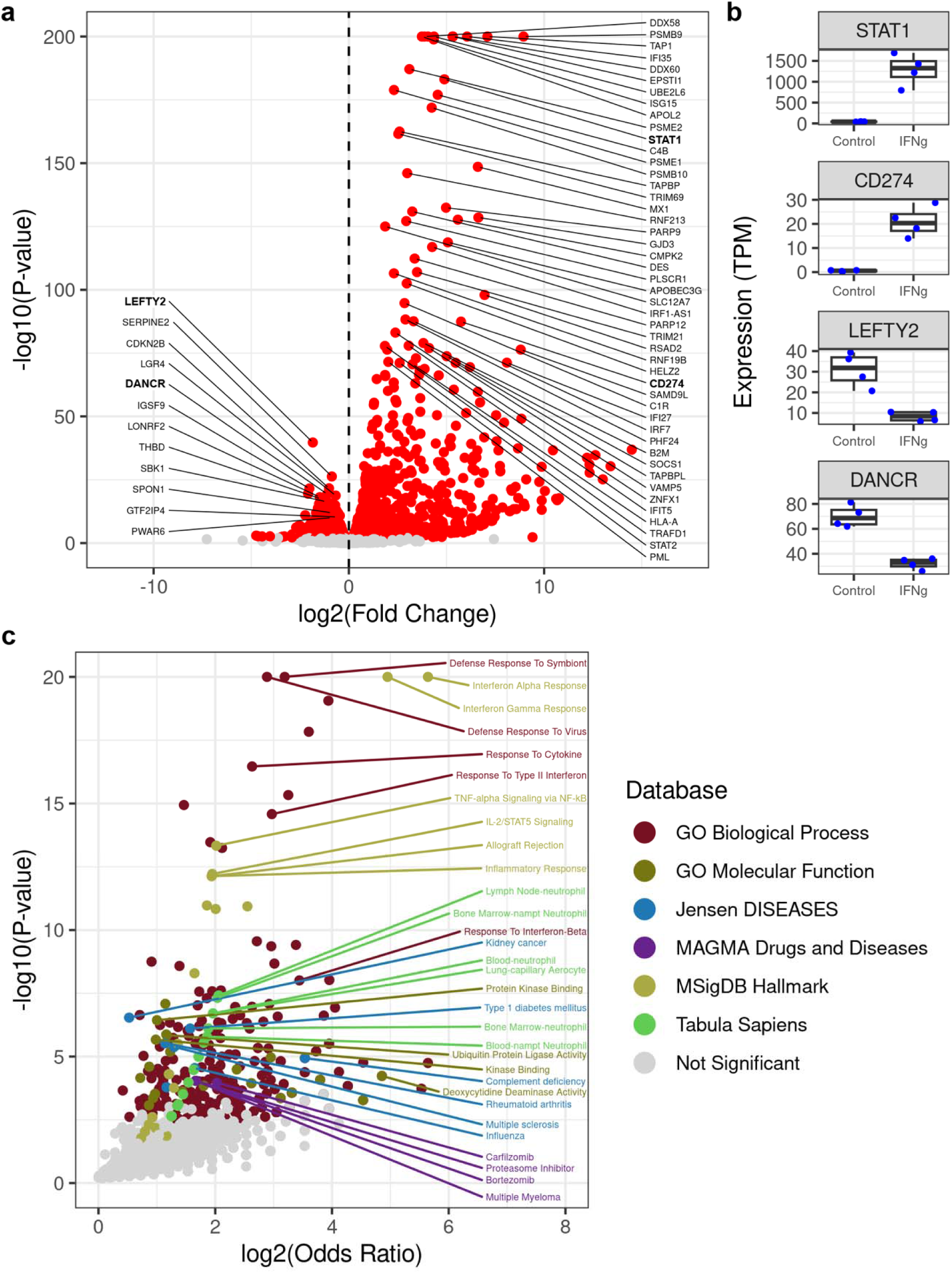
IFNγ stimulates an immune cell-independent inflammatory response. (a) Volcano plot showing 1,594 upregulated genes and 1,135 downregulated genes upon IFNγ treatment (n = 4 treated and n = 4 control). The x-axis is the log2 Fold Change, where positive values correspond to genes that are upregulated by IFNγ and negative values are downregulated genes, the y-axis is the - log10(P-value), and each point represents a gene. The points are colored by significance (FDR < 0.05), where “red” points are significant and “grey” points are not significant. The genes with bold labels are highlighted in panel b. (b) Box plots showing the expression of four differentially expressed genes; *STAT1* (first row), *CD274* (second row), *LEFTY2* (third row), and *DANCR* (fourth row). The x-axes contain the IFNγ treatment (Control and IFNg-treated), the y-axes contain the expression values (TPM), and the points correspond to the eight CVPC samples (n = 4 treated and n = 4 control). The line in the boxes represent the median, and the whiskers represent the 1.5x interquartile range (IQR). (c) Volcano plot showing functional enrichment of IFNγ upregulated genes in six Enrichr databases. The x-axis is the log2(Odds Ratio), the y-axis is the -log10(P-value), and each point represents a gene set. Points are colored by Enrichr database.

To identify biological processes affected by IFNγ stimulation, we performed functional enrichment analysis using Enrichr[26] on the 1,594 upregulated genes (Figure 1c, Table S3). In addition to the expected enrichments in immune-related terms, such as response to type 1 interferon (p = 1.3×10^-8^) and regulation of interleukin-12 production (p = 4.5×10^-7^), we observed enrichments in neutrophil gene signatures from different tissues, including lymph nodes, bone marrow, and blood from the *Tabula sapiens* gene set database [22] (Figure 1c). Several gene sets associated with autoimmune disorders (rheumatoid arthritis, multiple sclerosis, complement deficiency, and type 1 diabetes), cancers (kidney cancer and multiple myeloma), and cancer drugs (bortezomib and carfilzomib) were also enriched (Figure 1c).

Taken together, these results show that IFNγ stimulation in CVPCs upregulates expression of *CD274*, which encodes for PD-L1, consistent with what has been shown to occur in cardiac endothelial cells during inflammation [11,12]. Large expression differences in the JAK-STAT pathway and inflammatory response pathways also occur in the CVPCs notably in the absence of functioning immune cells.

### Cardiac endothelial cells drive the IFNγ stimulated inflammatory response

Given that *CD274* (PD-L1) expression is known to be upregulated in endothelial cells during inflammation[11,12], we hypothesized that upregulation of the inflammatory response pathways may also occur primarily in this cell type. To test this hypothesis, we obtained 180 iPSCORE fetal-like CVPCs, as well as 786 adult cardiovascular RNA-seq samples (left ventricle, atrial appendage, aorta, and coronary artery) from GTEx[21], performed a principal component analysis (PCA), and examined how the IFNγ treated samples were distinguished from their untreated matched samples in the PCA space (Figure 2a). As expected, PC1 discriminated between fetal-like CVPC and adult tissues, whereas PC2 separated adult cardiomyocyte (atrium and ventricle) from cardiac arterial (aorta and coronary artery) tissues. We observed that, while the transcriptomes of the four untreated samples from this study clustered with all the 180 iPSCORE CVPC samples, the transcriptomes of the treated samples shifted toward the GTEx cardiac arterial tissues. To confirm that the transcriptomes of the IFNγ treated CVPCs were becoming more similar to the transcriptomes of the GTEx cardiac arterial tissues compared to the transcriptomes of the GTEx cardiomyocyte-rich tissues, we collapsed the atrium and ventricle samples into a “CM” group and the aorta and coronary artery samples into an “Arterial” group and calculated the centroid coordinates. We then determined the Euclidean distances between the 8 CVPCs (treated and control) and the CM and Arterial centroids and performed a paired t-test to examine if IFNγ-treated CVPCs shifted toward the Arterial centroid. Indeed, we observed that the transcriptomes of the IFNγ-treated CVPCs shifted toward the arterial tissues (p-value = 4.6×10^-3^, data not shown), suggesting that IFNγ preferentially stimulates expression of genes in cardiac vascular tissue.

**Figure 2.**
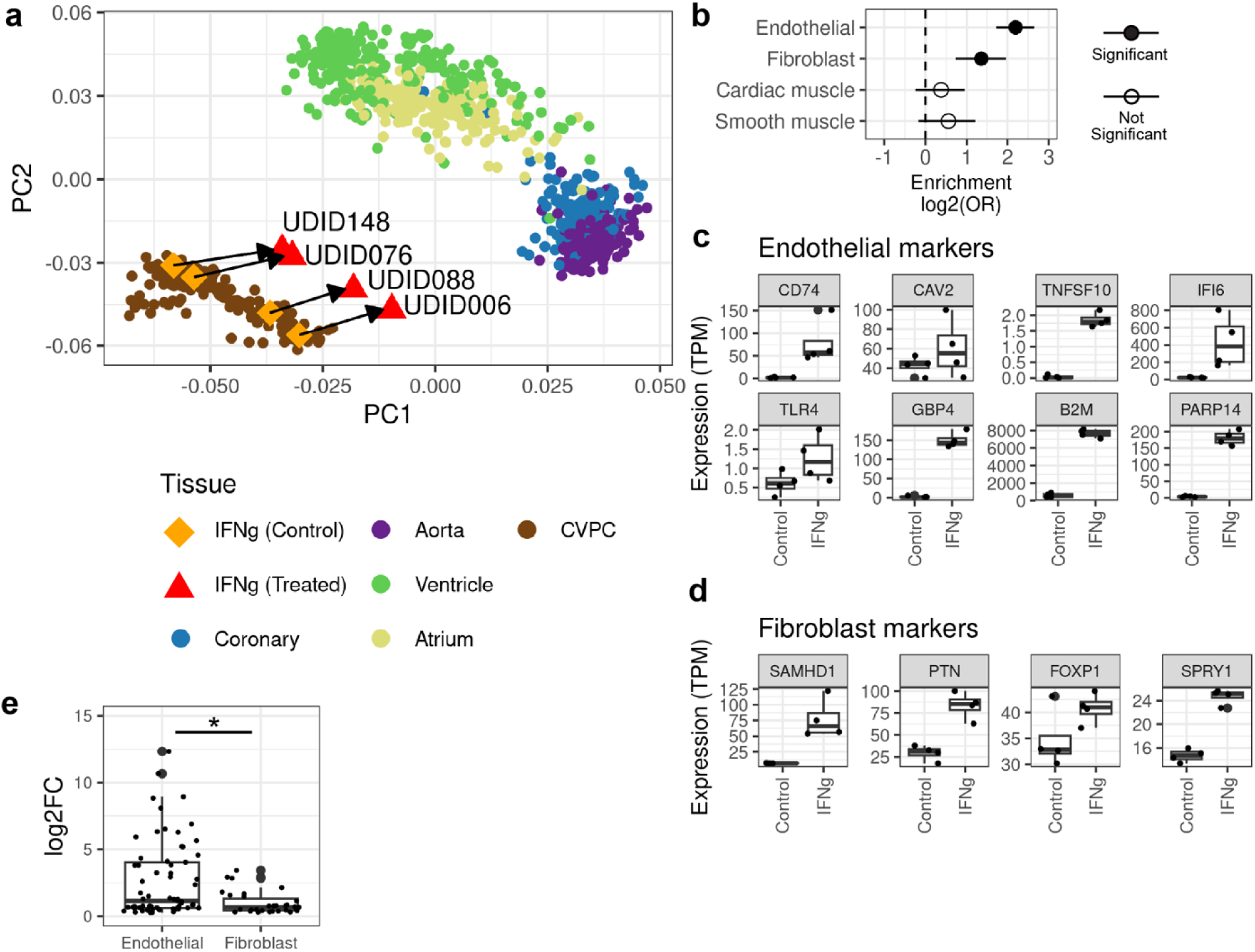
IFNγ-upregulated genes are enriched in endothelial markers. (a) PCA showing the clustering of the 180 iPSCORE CVPCs [14], 786 GTEx adult cardiac tissues[21], and 8 CVPC samples from this study. PCA was performed on the normalized expression of the 2,729 differentially expressed genes between control and treated samples. The 180 iPSCORE CVPCs and 786 GTEx adult cardiac tissues are represented by circles and are colored by tissue. The control CVPC controls from this study are represented by diamonds and colored orange, while the IFNγ-treated CVPCs are represented by triangles and colored red. The control (n = 4) and IFNγ-treated (n = 4) CVPC sample pairs are matched by their unique differentiation identifier (UDID). (b) Plot showing the enrichment of cell type marker genes in IFNγ-upregulated genes. Fisher’s Exact tests were performed to test the enrichment of cell type-specific marker genes in the set of IFNγ-upregulated genes, using the genes that were not differentially expressed by IFNγ treatment as background. The y-axis are the four cardiac cell types, and the x-axis is the log_2_(odds ratio). Solid points represent significant enrichments (Endothelial and Fibroblast) and hollow points represent non-significant enrichments (Cardiac muscle and Smooth muscle). The error bars represent the log2-transformed 95% confidence interval. (c-d) Box plots showing the expression of endothelial (c) and fibroblast (d) marker genes are upregulated by IFNγ. The y-axis is the gene expression (TPM) and the x-axis is the treatment (“Control” and “IFNg”). Each point represents a CVPC RNA-seq sample. The line in the boxes represent the median, and the whiskers represent the 1.5x interquartile range (IQR). e) Box plot comparing the differential expression of endothelial and fibroblast marker genes in IFNγ-treated CVPCs. The y-axis is the log_2_ log-fold change calculated by DESeq2 and the x-axis is divided by endothelial and fibroblast markers. The line in the boxes represent the median, and the whiskers represent the 1.5x interquartile range (IQR). The asterisk (*) indicates the difference between the two groups is significant by two-sided *t-test*.

To further examine the cell type-specificity of the response to IFNγ in CVPCs, we downloaded, filtered, and re-clustered single cell RNA-seq data on cardiac tissue obtained from 15 donors in the Tabula Sapiens dataset[22] (see Methods). After filtering, there were four clusters corresponding to cardiac muscle, endothelial, smooth muscle, and fibroblast populations (Figure S1a). Using Seurat[27], we next identified 784 genes that exhibited cell-type specificity, including 249 cardiac muscle, 209 endothelial, 167 fibroblast, and 162 smooth muscle-specific genes (Table S4). We performed Fisher’s Exact tests to evaluate whether the cell type-specific markers were enriched in IFNγ-upregulated genes. Consistent with our other observations, the endothelial-specific marker genes were the most enriched in the IFNγ-upregulated genes (Odds Ratio = 4.6, p-value = 9.8×10^-19^; Figure 2b). Fibroblast markers exhibited a weaker enrichment (Odds Ratio = 2.6, p-value = 2.5×10^-5^), while cardiac muscle and smooth muscle markers were not enriched (Figure 2b). We observed that endothelial markers that were upregulated by IFNγ have well characterized roles in immune responses, like *CD74*, *CAV2*, *TNFSF10, IFI6*, *TLR4*, *GBP4*, *B2M*, and *PARP14* (Figure 2c; Figure S2), while the fibroblast markers upregulated by IFNγ have roles in development and cell cycle regulation like *SAMHD1*, *PTN*, *FOXP1*, and *SPRY1* (Figure 2d) and generally exhibited more heterogeneous expression between the four matched control and treatment groups (Figure S3). Within the IFNγ-upregulated genes, the endothelial markers exhibited a significantly larger fold change (*t-test* p-value = 5×10^-5^) compared to the fibroblast (Figure 2e).

Taken together, these findings suggest that the IFNγ induced expression of the JAK-STAT and inflammatory response pathways primarily occurs in the cardiac endothelial cells, while fibroblasts may have a distinct non-immune response. Further investigation with a larger sample size or single cell data may be able to decipher how other cardiac cell types respond to IFNγ.

### IFNγ causes substantial chromatin remodeling

To profile the epigenomic landscape of cardiac cells treated with IFNγ, we performed ATAC-seq on the IFNγ treated CVPCs and their matched controls (Table S1) and identified 119,794 peaks. We performed differential chromatin accessibility analysis on IFNγ treated CVPCs and their paired controls and identified 3,603 differentially expressed peaks (FDR < 0.05), including 3,576 upregulated by IFNγ treatment and only 27 that were significantly downregulated (Table S5). These findings suggest that IFNγ activates a dormant regulatory network while not influencing networks that are active in homeostasis. To functionally characterize upregulated open chromatin, we performed gene set enrichment analysis using GREAT ontology[28], which annotates each peak with the Gene Ontology terms associated with its target genes. We found that the IFNγ-upregulated ATAC-seq peaks are associated with genes involved in interferon signaling, lymphocyte and mononuclear cell proliferation, and macroautophagy (Figure 3a, Table S6).

**Figure 3.**
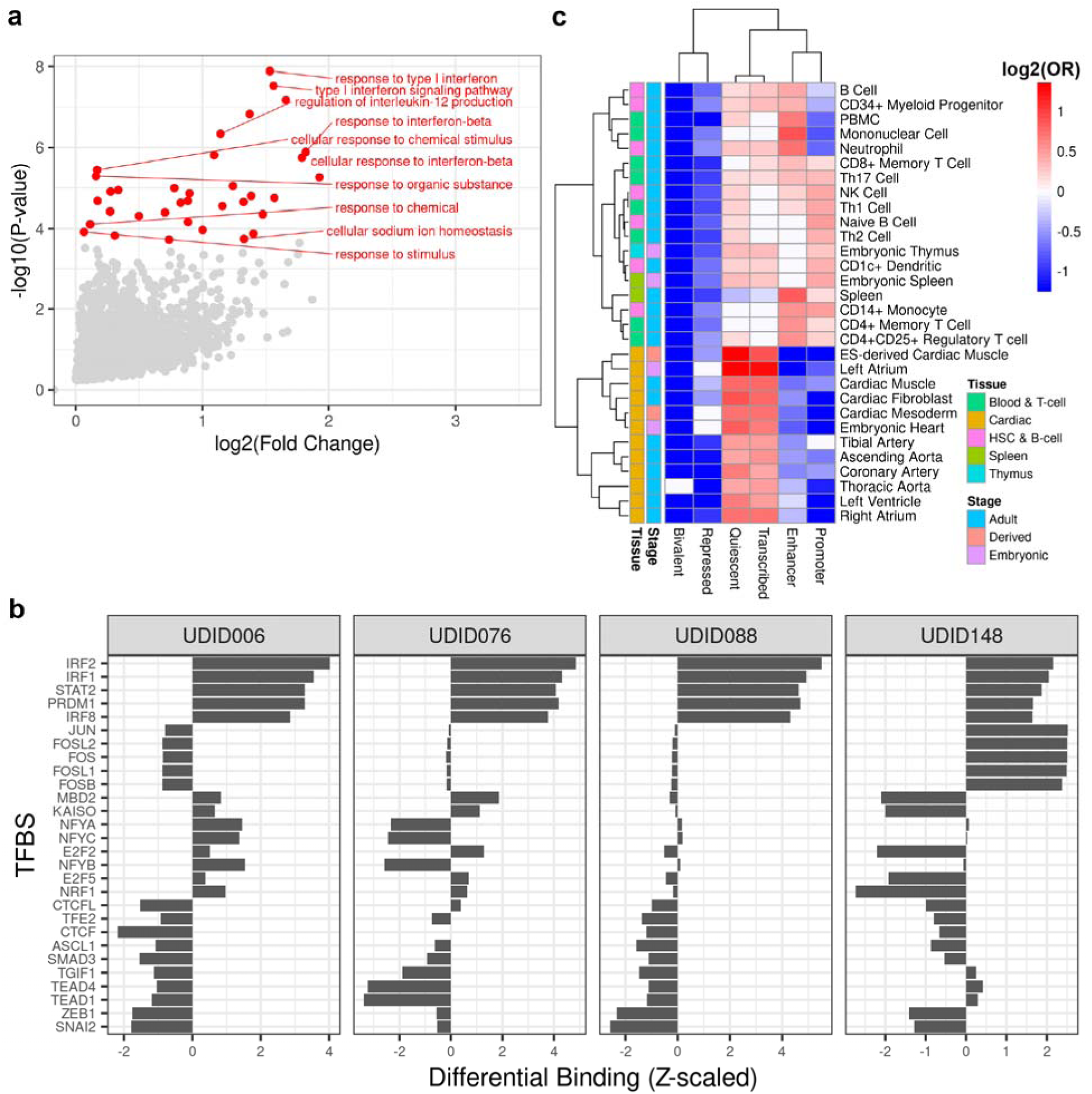
IFNγ activates an IRF and STAT regulatory network that is shared by immune cell types. (a) Volcano plot showing the enrichment in Gene Ontology gene sets associated with each upregulated peak using GREAT ontology. The x-axis is the log2(Fold Change), the y-axis is the -log10(P-value), and each point represents a gene set. Significant genes are colored red, while non-significant gene sets are colored grey. (b) Bar plots showing the differential binding of transcription factor binding sites in IFNγ-treated compared with control samples. The x-axis is the Z-scaled log2(Fold Change), the y-axis are the TFBSs that are among the top 10 most differentially bound in one or more of the CVPCs, and each panel corresponds to a CVPC differentiation (UDID) consisting of one set of paired IFNγ-treated and control samples. (c) Heatmap showing the differential enrichment of IFNγ upregulated open chromatin in cardiac and immune tissues. The x-axis contains the six collapsed chromatin states, the y-axis contains cardiac and immune tissues from the EpiMap repository, and the cells are filled with the log2(Odds Ratio) enrichments of 3,576 IFNγ upregulated open chromatin peaks compared to the 116,218 open chromatin peaks not upregulated by IFNγ treatment. The “Tissue” annotation column indicates whether the corresponding EpiMap tissue is immune (Blood & T-cell, HSC & B-cell, Spleen, Thymus) or cardiac. The “Stage” annotation column indicates whether the corresponding EpiMap tissue is Embryonic, Derived, or Adult.

Transcription factors (TFs) coordinate the expression of genes across the genome; therefore, we sought to identify which TFs were binding in IFNγ-upregulated chromatin. We performed TF footprinting analysis[29] using 401 TF motifs[30] in all 119,794 ATAC-seq peaks and evaluated differential TF binding between IFNγ-treated and control CVPCs (Figure 3b, Table S7). Consistent with the upregulation of genes in the JAK-STAT pathway, we observed that IRF1 and STAT2 TF binding sites (TFBSs) were differential bound across all IFNγ-treated CVPCs (Figure 3b). We also observed that developmental TEAD1 and SMAD3 TFBSs were differentially bound in three (UDID006, UDID076, and UDID088) of the four control CVPCs (Figure 3b). These results suggest that IFNγ treatment establishes an IRF/STAT-mediated regulatory network to activate the co-expression of inflammatory genes across the genome; and given the concordant IFNγ upregulation of genes in the JAK/STAT pathway (Figure 1a), suggests the open chromatin changes are primarily occurring in the endothelial cells.

### IFNγ establishes an immune regulatory network in cardiac endothelial cells

Since IRF and STAT family TFs are lowly expressed in cardiac tissue in homeostatic conditions (Figure 1a), we hypothesized that IFNγ upregulated chromatin that is normally inactive. We annotated the 119,794 ATAC-seq peaks with six chromatin states (Promoter, Enhancer, Transcribed regions, Bivalent, Repressed, and Quiescent) in 833 tissues from EpiMap[31]. For each tissue, we tested the enrichment of the 3,576 upregulated peaks in the six chromatin states, using the remaining 116,218 peaks that were not upregulated as background (Figure 3c, Table S8). We observed that upregulated peaks were enriched in quiescent chromatin and transcribed regions and strongly depleted in promoters and enhancers in cardiac tissue (Figure 3c), supporting that IFNγ opens chromatin that is not accessible in homeostatic cardiac conditions. Additionally, we found that IFNγ upregulated peaks were most strongly enriched in immune cell type promoters and enhancers (Figure 3c), suggesting that the IFNγ induced regulatory network in the cardiac endothelium is shared with immune cell types.

Taken together, our results indicate that IFNγ modulates dramatic chromatin remodeling in the cardiac endothelium by activating an immune cell-like, IRF/STAT-mediated regulatory networks.

### Cardiac disease-associated variants overlap IFNγ-upregulated ATAC-seq peaks

To evaluate cardiac disease-associated variants that overlap CVPC open chromatin, we first downloaded genome-wide association study (GWAS) summary statistics for cardiac diseases, including cardiac dysrhythmias, atrioventricular and left bundle-branch block, coronary atherosclerosis, hypertension, angina pectoris, acute myocardial infarction, chronic ischemic heart disease, atrial fibrillation, and heart failure. We then extracted 28,543 cardiac disease-associated variants (p-value < 5×10^-8^, minor allele frequency ≥ 1%) from the summary statistics and determined their overlap with the 119,794 CVPC ATAC-seq peaks. In total, 652 unique disease-associated variants were in 444 CVPC ATAC-seq peaks (Figure 4a). Of these 652 variants, 29.4% (n=192) were associated with multiple cardiac diseases.

**Figure 4.**
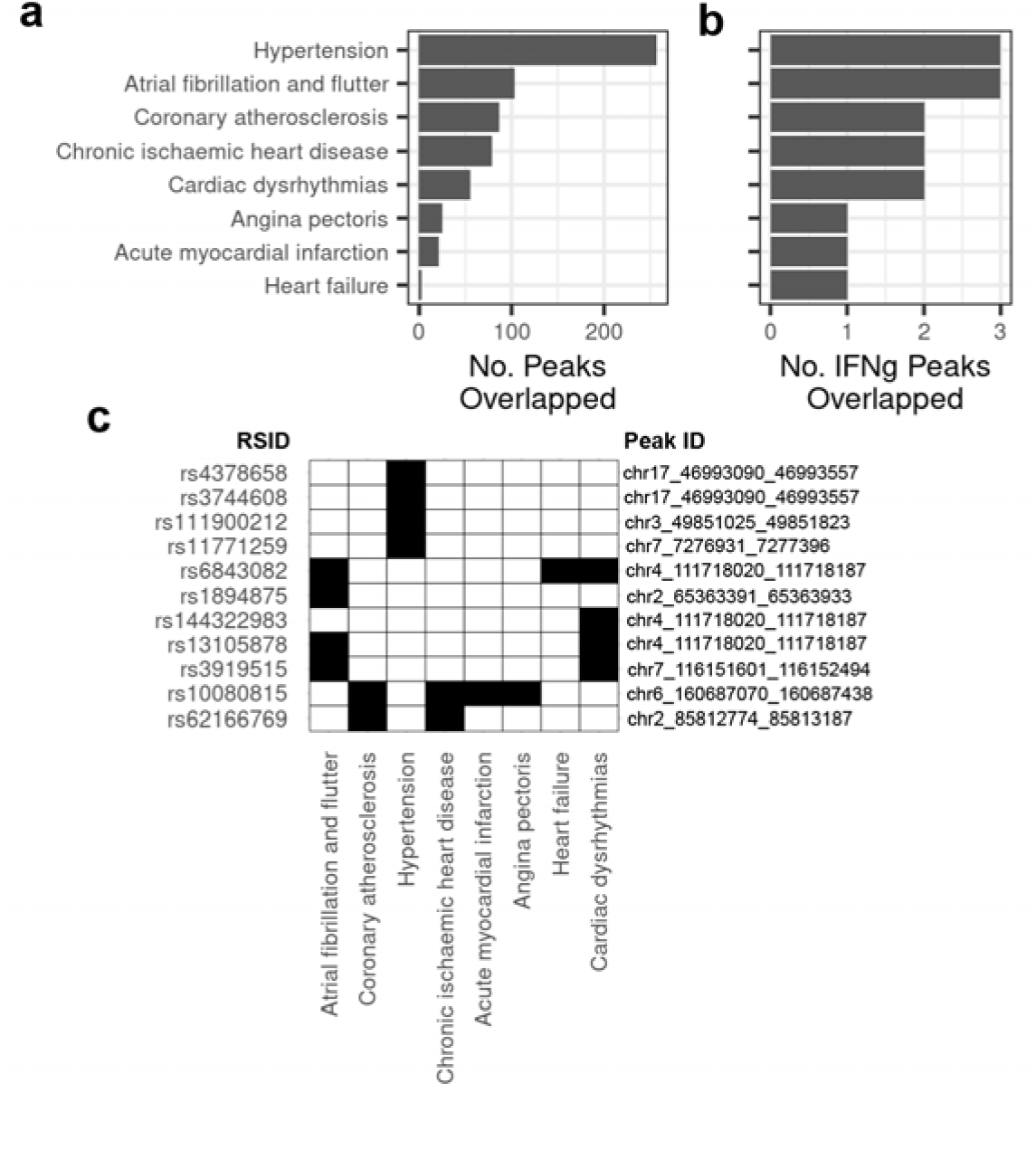
Cardiac disease associated variants are in IFNγ-upregulated ATAC-seq peaks. a-b) Bar plots showing (a) the total number of CVPC and (b) IFNγ-upregulated ATAC-seq peaks that overlap cardiac disease associated variants. The y-axis shows the cardiac disease GWAS and the x-axis is the number of unique (a) CVPC and (b) IFNγ-upregulated ATAC-seq peaks that overlapped at least one disease-associated variant. The y-axis is shared for both panels. c) Binary heatmap showing cardiac disease-associated variants in IFNγ-upregulated ATAC-seq peaks. The left y-axis contains the rsids of the cardiac disease-associated variants, the right y-axis are the chromosomal positions of the IFNγ-upregulated peaks (Table S9) that contain at least one the cardiac disease-associated variant, the x-axis contains the eight cardiac diseases with at least one variant in an in IFNγ-upregulated peak, and the cells are filled “black” to indicate that the variant is associated with the corresponding disease.

To assess whether certain GWAS signals are explained by regulatory variation that is only active in cardiac tissues after IFNγ stimulation, we focused on 11 unique disease-associated variants in 8 IFNγ-upregulated peaks (Figure 4b). Of these 11 disease-associated variants, 5 were associated with multiple traits (Figure 4c, Table S9). Interestingly, rs10080815 (chr6:160687412:T>G) overlaps an IFNγ-upregulated peak and is shared in four cardiac diseases (angina pectoris, myocardial infarction, chronic ischemic heart disease, and coronary atherosclerosis). rs10080815 is in an adult aorta expression quantitative trait locus (eQTL) for *SLC22A1*[21], which encodes an organic cation transporter, that has previously been associated with coronary artery disease, cardiovascular disease, and blood lipid levels[32]. rs6843082 (chr4:111718067:G>A) is also in an IFNγ-upregulated peak and is significant in GWAS for atrial fibrillation and flutter, cardiac dysrhythmias, and heart failure (Figure 4c). rs6843082 is located ∼150kb upstream of *PITX2*, which is encodes a transcription factor involved in myocardial development[33].

Taken together, these findings provide evidence for context-specificity of GWAS loci by showing that certain cardiac disease variants are in chromatin that is only open during inflammation. These analyses can serve as a proof of-principle and justification for generating more data for larger scale studies.

## Discussion

iPSC-derived tissues are a powerful model for assessing the effects of chemical or environmental stimuli on human tissues that are difficult to obtain. In this study, we demonstrate the utility of using iPSC-derived cardiovascular progenitor cells (CVPCs) to model the effects of a pro-inflammatory cytokine, IFNγ, on human cardiac tissue. We performed RNA-seq and ATAC-seq on four paired IFNγ-treated and control CVPCs to characterize the transcriptomic and epigenomic changes. Despite the absence of immune cell types, IFNγ stimulated the upregulation of *CD274* (PD-L1) and JAK-STAT-related genes and mounted an immune-like response in cardiac tissue (Figure 1). Additionally, IFNγ stimulation downregulated key developmental genes (*LEFTY1*, *DANCR*). Indeed, by integrating RNA-seq from 786 adult cardiac samples from four tissues (aorta, atrium, coronary artery, and left ventricle) and 180 iPSCORE CVPCs, we observed that transcriptomes of IFNγ-stimulated CVPCs shifted toward the aorta and coronary samples relative to their paired controls in the PC space (Figure 2a). Expression analysis of cell type-specific genes revealed that these IFNγ-induced transcriptomic changes primarily occurred in the cardiac endothelial cell population (Figure 2b-c). While we observed weak evidence for an independent fibroblast response to IFNγ, the small sample size limited our ability to further characterize it. Our study is consistent with previous observations that demonstrate that the vascular endothelium mediates a barrier response to pro-inflammatory cytokines in the heart[11,12].

Differential chromatin accessibility and TF footprinting analyses revealed that IFNγ stimulation also upregulated regulatory elements bound by IRF and STAT family TFs that were near genes in immune pathways (Figure 3a-b). We show that regulatory elements upregulated by IFNγ are inactive in several cardiac tissues but are active enhancers in dozens of immune cell types, suggesting that the cardiac endothelium adopts an immune-like phenotype during IFNγ induced inflammation. We also show that GWAS variants for 8 cardiac diseases overlap ATAC-seq peaks that are only present after IFNγ treatment, suggesting that they may capture context-specific regulatory variants. Our findings can be used to inform experimental and animal studies to study the mechanisms of immune-related cardiac disease. Future studies can perform quantitative trait loci analyses on gene expression and chromatin accessibility using hundreds of IFNγ-treated CVPCs to identify genetic risk factors associated with immune-related cardiac diseases and phenotypes. These larger scale studies would also be better powered to discover additional cell type-specific responses to IFNγ treatment that we did not identify due to small sample size.

In summary, our results provide biological insights into cardiac tissue cell type-specific responses to IFNγ stimulation and demonstrate the utility of using iPSC-derived tissues to characterize drug exposure and other context specific contributions to disease.

## Code Availability

Relevant scripts and notebooks are publicly available at https://github.com/frazer-lab/EndotheliumIFNg.

## Data Availability

FASTQ sequencing data for 8 RNA-seq and 8 ATAC-seq samples have been deposited into GSE263611. TOBIAS TF binding predictions, gene expression matrix, signature gene matrix for deconvolution, the ATAC-seq peaks and summits, and the GREAT ontology output results have been deposited in Figshare: (https://figshare.com/projects/Endothelium_IFNg/202329).

## Contributions

TDA, MD, and KAF conceived the study. TDA, INJ, JPN, and iPSCORE consortium members performed the computational analyses. MD and KAF oversaw the study. ADC and members of the iPSCORE Consortium performed the differentiations and generated molecular data. TDA, INJ, MD, and KAF prepared the manuscript.

## Acknowledgments

This work was supported by the National Library Training Grant T15LM011271, the National Institute of Diabetes and Digestive and Kidney Disease (NIDDK) F31DK131867 and P30DK063491, the National Heart, Lung and Blood Institute (NHLBI) F31HL158198 and U01HL107442, and the National Human Genome Research Institute (NHGRI) RM1HG011558. Additional support was also received from a California Institute for Regenerative Medicine grants GC1R-06673-B and EDUC2-08388, and NSF-CMMI division award 1728497.

## Artificial Intelligence Disclosure

The authors did not use generative AI or AI-assisted technologies in the development of this manuscript.

## SUPPLEMENTAL FIGURES

**Figure S1.**
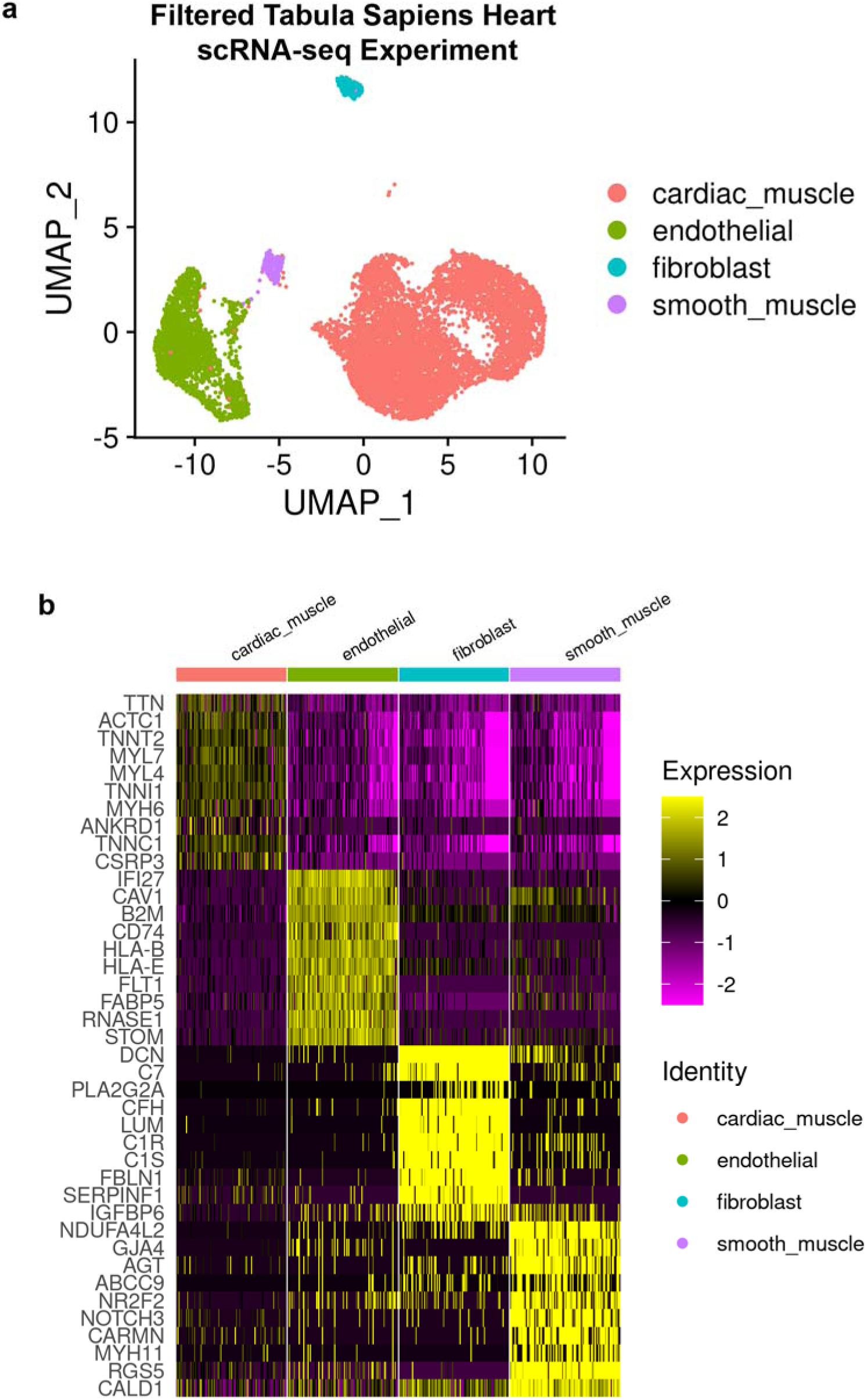
Identification of cardiac cell type markers using the Tabula Sapiens. a) UMAP of the filtered single cell RNA-seq heart dataset from the Tabula Sapiens project. The axes correspond the UMAP coordinates, and each point represents a cardiac cell. The colors correspond to the cell types that are assigned to the cells. b) Heatmap showing the expression of the top 10 marker genes for the four cell types. The y-axis contains the 40 marker genes, the x-axis are a random subset of 100 cells from each cell type, the cells are filled with the expression of the marker genes.

**Figure S2.**
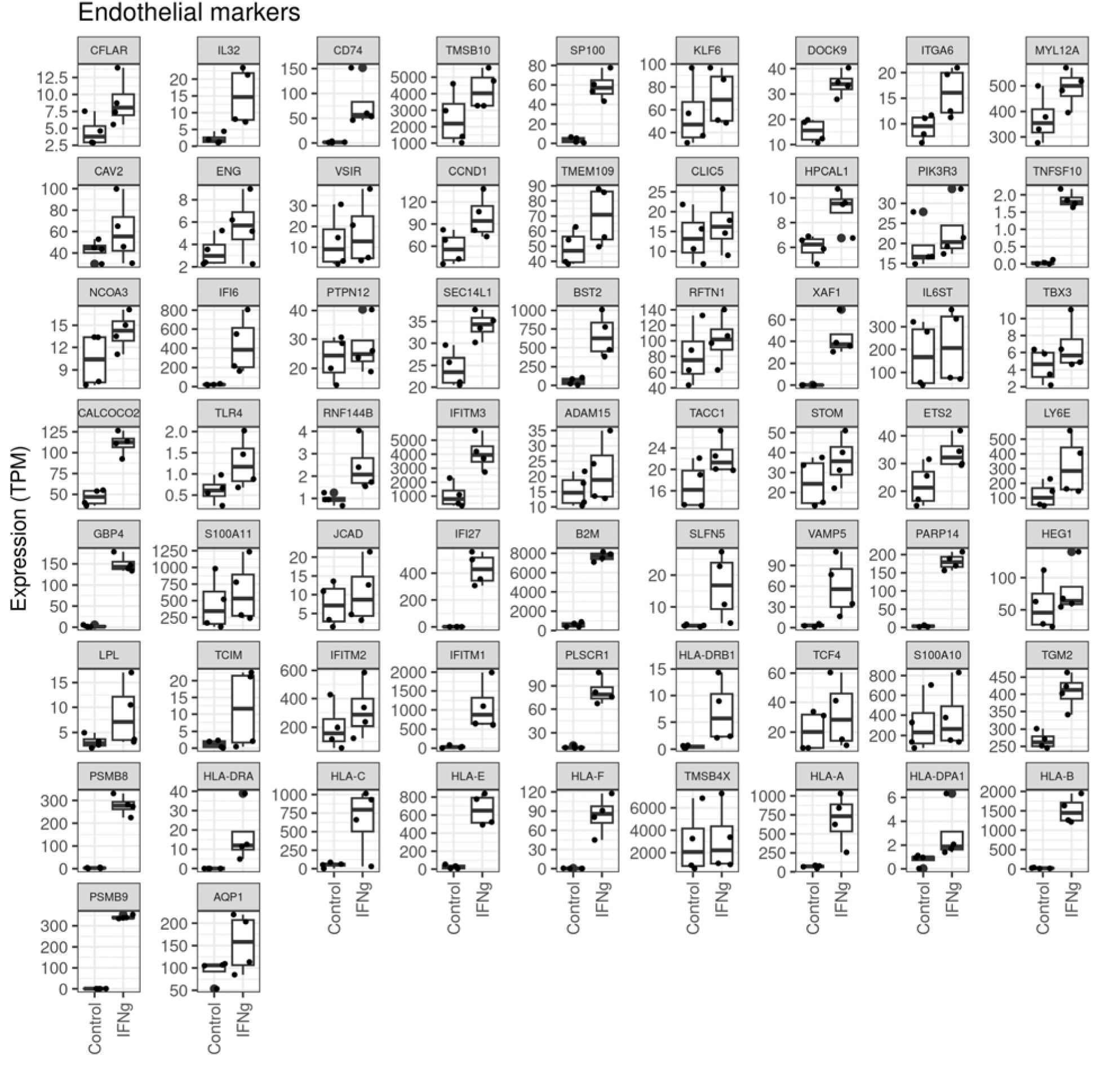
Expression of the IFNγ-upregulated endothelial markers in treated and control CVPCs. Box plots showing the expression of IFNγ-upregulated endothelial markers in treated and control CVPCs. The x-axis is split by the treatment groups (“Control” and “IFNg”) and the y-axis is the gene expression (TPM). Each point represents a CVPC RNA-seq sample. The line in the boxes represent the median, and the whiskers represent the 1.5x interquartile range (IQR).

**Figure S3.**
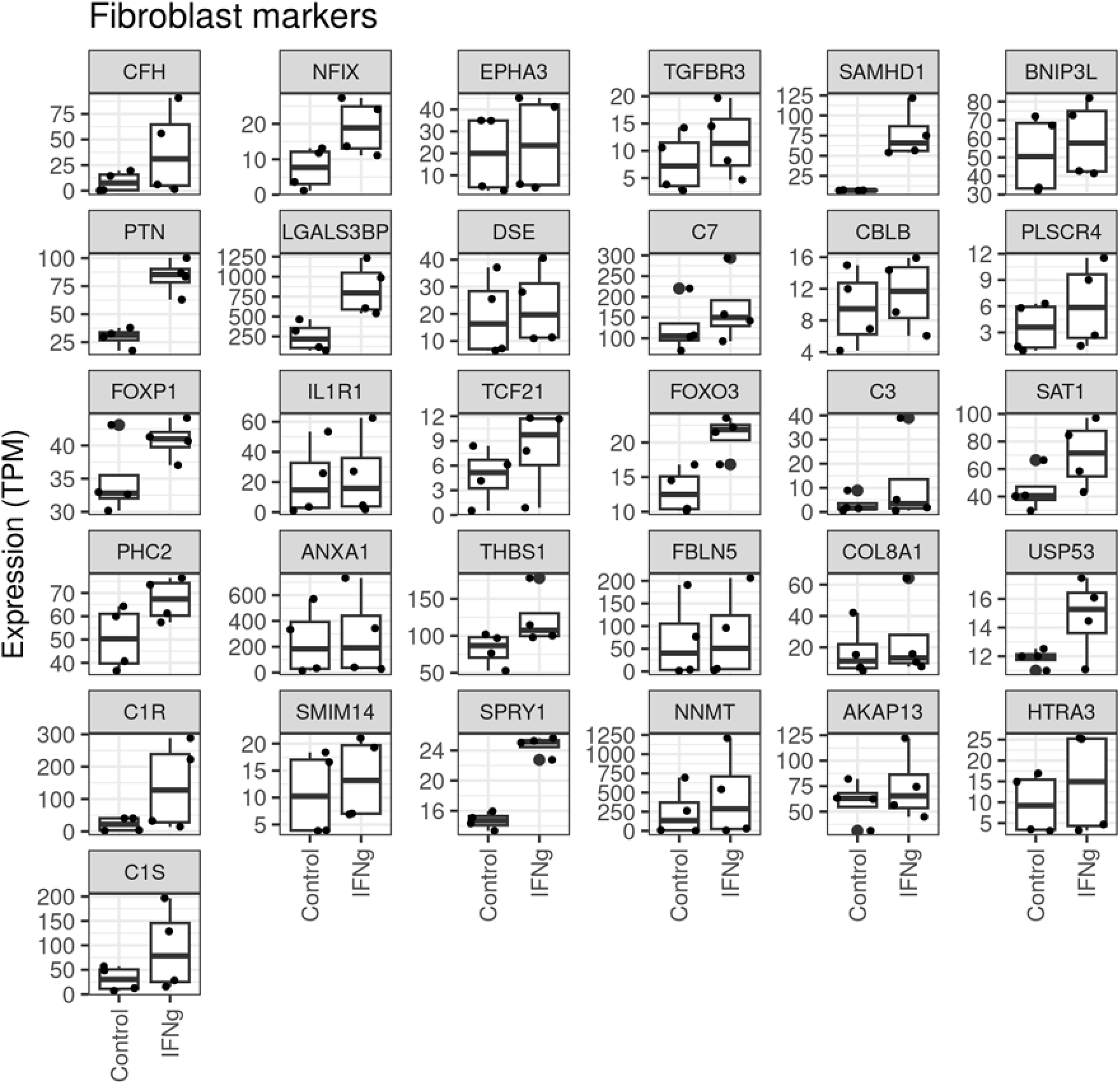
Expression of the IFNγ-upregulated fibroblast markers in treated and control CVPCs. Box plots showing the expression of IFNγ-upregulated fibroblast markers in treated and control CVPCs. The x-axis is split by the treatment groups (“Control” and “IFNg”) and the y-axis is the gene expression (TPM). Each point represents a CVPC RNA-seq sample. The line in the boxes represent the median, and the whiskers represent the 1.5x interquartile range (IQR).

## METHODS

### 1. CVPC culture and treatment conditions

iPSC lines from four unrelated individuals recruited as part of the iPSCORE project were previously differentiated into CVPCs using small molecules and enriched using metabolic selection with lactate[13,14] (Table S1). The CVPCs were harvested at day 25, which we and others have previously shown represent fetal cardiac cells[34], and cryopreserved at -80° C. The cryopreserved CVPCs (day 25) were then thawed, plated at a density of 2.5×10^5^/cm^2^ cells on Matrigel (Corning) coated 6-well plates, and allowed 72 hours for recovery and expansion. On day 28 (three days after thawing), CVPCs were treated with 100 nM human recombinant IFNγ (ThermoFisher) for 24h. On day 29, control and IFNγ-treated CVPC samples (eight in total) were harvested. Each iPSC differentiation into CVPCs was given a unique differentiation identifier (UDID). The four control and four IFNγ-treated CVPC samples are matched by their UDID.

### 2. Molecular data generation and sequencing

#### 2.1 RNA-seq

We extracted total RNA from lysed pellets frozen in RLT Plus buffer (collected on day 29) using the Quick-RNA™ MiniPrep Kit (Zymo Research), assessed quality to make sure RNA Integrity number (RIN) was 7.5 or greater, prepared indexed libraries using the Illumina TruSeq stranded mRNA kit and sequenced with 150 bp paired-end reads on an Illumina HiSeq 4000 (mean = 91.4M reads).

#### 2.2 ATAC-seq

ATAC-seq samples were processed as previously described in detail[25]. Briefly, frozen nuclear pellets of 1×10^5^ CVPC were thawed on ice and tagmented in total volume of 25 μl in permeabilization buffer containing digitonin (10 mM Tris-HCl pH 7.5, 10 mM NaCl, 3 mM MgCl^2^, 0.01% digitonin) and 2.5 μl of Tn5 from Nextera DNA Library Preparation Kit (Illumina) for 45-75min at 37°C in a thermomixer (500 RPM shaking). To eliminate confounding effects due to index hopping, all libraries within a pool were indexed with unique pairs of i7 and i5 barcodes. Libraries were amplified for 12 cycles using NEBNext^®^ High-Fidelity 2X PCR Master Mix (NEB) in total volume of 25µl in the presence of 800 nM of barcoded primers (400nM each) custom synthesized by Integrated DNA Technologies (IDT) and sequenced with 150-bp paired end reads on a HiSeq4000 (mean = 112.9M reads).

### 3. Data Processing

#### 3.1 RNA-seq

To measure gene expression in the 8 CVPC samples, we obtained raw counts in transcript per million bp (TPM) as previously described[15]. Briefly, FASTQ files were aligned to the hg19 reference human genome using with STAR v.2.7.3[35] and GencodeV.34lift37 with parameters: *-- outFilterMultimapNmax 20 --outFilterMismatchNmax 999 --alignIntronMin 20 --alignIntronMax 1000000 --alignMatesGapMax 1000000*. Next, the BAM files were sorted and indexed using sambamba v0.6.7[36], and duplicates were marked using bammarkduplicates from the biobambam software[37]. To quantify gene expression (TPM), we used RSEM version 1.2.20[38] with the following parameters: *rsem-calculate-expression –bam --estimate-rspd --paired-end --forward-prob 0*. RNA-seq sample quality was analyzed with samtools[39] stats, idxstats, and Picard RNA Metrics (Table S1).

#### 3.2 ATAC-seq

The 16 FASTQ files were aligned to the hg19 reference genome with STAR using the following parameters; *--outFilterMultimapNmax 20, -- outFilterMismatchNmax 999, -- outFilterMismatchNoverLmax 0.04, --seedSearchStartLmax 20, -- outFilterScoreMinOverLread 0.1, outFilterMatchNminOverLread 0.1*. Sample mate coordinates were filled using samtools fixmate and duplicates were marked using samtools markdup[39]. We then removed poorly mapped reads, fragments < 38 bp, fragments > 2,000 bp, and reads that did not map to autosomal or sex chromosomes with samtools view[39]. Finally, we removed reads in the blacklisted regions (https://mitra.stanford.edu/kundaje/akundaje/release/blacklists/hg19-human/wgEncodeHg19ConsensusSignalArtifactRegions.bed.gz) using bedtools intersect[40] with the *v* parameter.

### 4. Analysis

#### 4.1 RNA-seq

##### 4.1.1 Differential Gene Expression

The standard DESeq2 (version 1.34.0)[41] workflow for identifying differentially expressed genes was followed. Briefly, the counts and associated metadata for all 8 RNA-seq samples were loaded into a DESeq2 object using the *DESeqDataSetFromMatrix* function. To account for cell type differences between samples, the design was specified as ∼ UDID + treatment and the *DESeq* function was applied to identify genes that are differentially expressed (Benjamini-Hochberg’s Adjusted P-value < 0.05) by IFNγ treatment.

##### 4.1.2 Functional Enrichment Analysis

The 1,594 upregulated genes DESeq2 were used as input into Enrichr[26] and functional enrichment analysis was performed with standard parameters on all libraries cumulatively containing >420k gene sets. For plot legibility, the following gene set libraries were prioritized; Jensen DISEASES, MAGMA Drugs and Diseases, Tabula Sapiens, GO Biological Process 2023, GO Molecular Function 2023, MSigDB Hallmark 2020, and IDG Drug Targets 2022.

##### 4.1.3 Integrated PCA

We obtained gene expression (TPM) values for 180 iPSCORE CVPC samples, 207 adult left ventricle, 187 atrial appendage, 215 aorta, and 117 coronary artery samples from GTEx using processed data from our previous studies (10.6084/m9.figshare.16920658)[15,21], combined these with the TPM values for the IFNγ-treated and control samples, and performed quantile normalization using the *normalize.quantiles* function from the preprocessCore package and qnorm functions in R, and obtained mean expression = 0 and standard deviation = 1 for each gene. We extracted the normalized expression levels of the 2,729 differentially expressed genes (1,594 upregulated and 1,135 downregulated) and performed PCA using the prcomp function in R.

##### 4.1.4 Determining cardiac cell type markers

We obtained an annotated scRNA-seq Seurat h5ad object from the *Tabula sapiens* heart dataset (https://figshare.com/articles/dataset/Tabula_Sapiens_release_1_0/14267219) which is composed of 11,505 cells annotated with six cell types (cardiac muscle, smooth muscle, endothelial, fibroblasts, macrophages, and hepatocytes). We converted the annotated Seurat h5ad object to an h5seurat file, using the *Convert* function from the SeuratDisk R package. We loaded the file into Seurat (version 4.3.0)[27] and removed cells annotated as “macrophages” and “hepatocytes” because the lactate selection step in the CVPC differentiation results in survival of only cardiac lineage cell types. After filtering, we renormalized, rescaled, and reclustered the cells at resolution = 0.08, using Seurat. We annotated 5 of the 6 resulting clusters, using original *Tabula sapiens* annotations. Two clusters mapped to cardiac muscle, one cluster mapped to smooth muscle, endothelial, and fibroblast each, yielding four cell type clusters. The unassigned cluster exhibited increased expression of markers for antigen presenting cells and was composed of < 60 cells, therefore we removed it. We then ran the Seurat FindAllMarkers function using the parameters *logfc.threshold = 0.1, only.pos = TRUE,* and *min.pct = 0.2* to obtain marker genes and classify cardiac cell types belonging to the four remaining cardiac cell types in *Tabula sapiens*.

##### 4.1.5 Calculating the enrichment of cell type markers in IFNγ-upregulated genes

Using the cell type-specific genes identified from the *Tabula Sapiens* scRNA-seq heart dataset (see section: 4.1.4 Determining cardiac cell type markers), we determined the overlap of the marker genes and the IFNγ-upregulated genes. First, we removed all genes that were not expressed in the control and IFNγ-treated CVPCs. We next removed all genes that were downregulated by IFNγ treatment to obtain genes that are not differentially expressed after IFNγ treatment. For each of the four cell types, we performed a two-sided Fisher’s Exact test to test the enrichment of cell type-specific genes in IFNγ-upregulated genes, using the genes that did not exhibit differential expression after IFNγ treatment as background.

#### 4.2 ATAC-seq

##### 4.2.1 MACS2 Peak Calling

Narrow peaks were called using MACS2[42] on each of the 8 BAM files individually, using the following parameters; -f BAMPE -g hs --nomodel --shift -100 --extsize 200 --call-summits –q 0.01, which resulted in an average of 77,643 ± 33,040 peaks per sample. We merged the peak bed files from the 8 ATAC-seq samples, then removed peaks that were only present in one sample which resulted in 119,794 reference peaks. For each sample, we counted the number of reads in the reference peaks using featureCounts[43].

##### 4.2.2 Differential Chromatin Accessibility

The standard DESeq2 (version 1.34.0)[41] workflow for identifying differentially accessible peaks was followed. Briefly, the counts and associated metadata for all 8 ATAC-seq samples were loaded into a DESeq2 object using the DESeqDataSetFromMatrix function. To account for cell type differences between the four control (untreated) CVPC samples, the design was specified as ∼ UDID + treatment and the DESeq function was applied to identify ATAC-seq peaks that are differentially accessible (Benjamini-Hochberg’s Adjusted P-value < 0.05) by IFNγ treatment.

##### 4.2.3 GREAT ontology functional enrichment of upregulated peaks

The 3,576 upregulated peaks from DESeq2 were used as input, and all 119,794 peaks as background, to perform functional enrichment analysis on Gene Ontology Biological Process, Molecular Function, and Cellular Component gene sets using GREAT[28].

##### 4.2.4 Transcription Factor Binding Site Predictions

We used TOBIAS[29] to predict binding motifs and map potential transcription factor occupancy sites across our peaks to profile chromatin accessibility after IFNγ stimulation. We ran TOBIAS ATACorrect on the 8 BAM files to correct for cut site biases introduced by the Tn5 transposase within the 119,794 ATAC-seq peaks using the following parameters: --genome hg19 fasta and – blacklist hg19blacklist.v2.bed (from https://mitra.stanford.edu/kundaje/akundaje/release/blacklists/hg19-human/wgEncodeHg19ConsensusSignalArtifactRegions.bed.gz). Next, we calculated footprint scores with TOBIAS ScoreBigWig, using the BED file containing the coordinates of all the 119,794 ATAC-seq peaks. Finally, to calculate differential transcription factor binding between control and IFNγ treated CVPCs, we ran TOBIAS BINDetect on each of the four control and IFNγ-treated CVPC samples matched by their UDIDs using position weight matrices (PWM) for 401 HOCOMOCO V.11[30] transcription factors, using the following parameters: --motif *HOCOMOCO PWM file* –signals *untreatedUDID.bw treatedUDID.bw*. For plot legibility, we plotted the average fold change and p-value across the four UDIDs for 265 TFs with motif quality (“A” or “B”).

##### 4.2.5 Differential Chromatin State Enrichment

We obtained 18 ChromHMM chromatin states from 833 tissues from the Epimap Repository[31] (https://personal.broadinstitute.org/cboix/epimap/ChromHMM/observed_aux_18_hg19/CALLS/). For all 833 tissues, we collapsed the 18 chromatin states into 6 states, including Promoter (“TssA”, “TssFlnk”, “TssFlnkU”, “TssFlnkD”), Enhancer (“EnhA1”, “EnhA2”, “EnhG1”, “EnhG2”, “EnhWk”), Transcribed (“Tx”, “TxWk”), Bivalent (“TssBiv”, “EnhBiv”), Repressed (“ReprPC”, “ReprPCWk”, “ZNF/Rpts”, “Het”), and Quiescent (“Quies”). We used the coordinates of the maximum summit of each peak to annotate all 119,794 ATAC-seq peaks with the 6 collapsed chromatin states from all 833 EpiMap tissues, using bedtools intersect[40]. For each tissue, we performed Fisher’s Exact tests to test the differential enrichment of 3,576 IFNγ-upregulated ATAC-seq peaks in the six chromatin states using the 116,218 ATAC-seq not upregulated by IFNγ treatment as background.

##### 4.2.6 Identifying GWAS variants located in IFNγ-upregulated ATAC-seq peaks

We downloaded GWAS for 9 cardiac diseases (angina pectoris, acute myocardial infarction, chronic ischaemic heart disease, atrioventricular and left bundle-branch block, atrial fibrillation and flutter, heart failure, hypertension, coronary atherosclerosis, and cardiac dysrhythmias) from the Pan-UK Biobank summary statistics repository (https://pan-ukb-us-east-1.s3.amazonaws.com/sumstats_flat_files/; Table S9). From each summary statistic file, we filtered variants with the low allele frequencies (case allele frequency < 1%) and non-genomewide significant meta-analysis p values (p-value > 5×10^-8^), yielding 28,543 cardiac disease-associated variants. We then used *bedtools intersect* [40] to identify cardiac disease-associated variants that overlap any of the CVPC 119,794 ATAC-seq peaks, then focused on the 3,576 IFNγ-upregulated ATAC-seq peaks.

## SUPPLEMENTAL TABLE LEGENDS

**Table S1: Study Metadata**

Sheet 1: Subject Metadata: This table contains information on the iPSCORE subject whose data was analyzed in this study. The columns describe the **iPSCORE_ID** and the **Subject_UUID** which is a universal unique identifier, the iPSCORE **Family_ID**, the **Sex** of the donor, and the **Age_at_Enrollment**.

Sheet 2: RNA-seq Sample Metadata: This table contains information and quality metrics of the RNA-seq samples analyzed in this study. The columns describe the **UDID** the CVPC unique differentiation identifier, **Treatment** either untreated “CONTROL” or “IFNg” treated, **Sample_UUID** the sample universal unique identifier, the **iPSCORE_ID** and **Subject_UUID** of the donor, and the quality control metrics of the samples, including the number of **Total Reads**, **Duplicates**, **Mapped** reads, and **Properly Paired** reads.

Sheet 3: ATAC-seq Sample Metadata: This table contains information and quality metrics of the RNA-seq samples analyzed in this study. The columns describe the **UDID** the CVPC unique differentiation identifier, **Treatment** either untreated “CONTROL” or “IFNg” treated, **Sample_UUID** the sample universal unique identifier, the **iPSCORE_ID** and **Subject_UUID** of the donor, and the quality control metrics of the samples, including the number of **Total Reads**, **Mapped** reads, and **Properly Paired** reads.

**Table S2: Differential Gene Expression**

This table contains the output from differential gene expression analysis performed using the DESeq2 R package. The columns include the **Gene Name** and **Gene ID** from Gencode.v34lift37, and the standard output of DESeq2, including **baseMean**, **log2FoldChange**, **lfcSE** (standard error of the log2 fold change), **stat** (the Wald statistic), **p-value**, and the Benjamini Hochberg **Adjusted p-value**. Non-significant (adjusted p-value > 0.05), IFN-γ upregulated (adjusted p-value < 0.05 & log2FoldChange > 0) and IFN-γ downregulated (adjusted p-value < 0.05 & log2FoldChange < 0) genes are reported.

**Table S3: Enrichr Gene Set Enrichment Results**

This table contains the output for the gene set enrichment analysis performed using the Enrichr R package. The columns describe the **Database** in Enrichr, the **Term** for the gene set, the **Number of Upregulated Genes in the Term**, the **Total Number of Genes in the Term**, the nominal **P-value**, Benjamini Hochberg **Adjusted P-value**, and the **Odds Ratio**.

**Table S4: Cell Type-Specific Markers**

This table includes cell type-specific marker genes identified using the *Tabula Sapiens* scRNA-seq heart dataset. It contains columns describing the **Cell_Type** (Cardiac Muscle, Smooth Muscle, Endothelial, or Fibroblast), **Gene Name**, the average log_2_ fold-change (**Average_Log2FC**), **P_value**, and the **Adjusted_P_value** calculated the *FindAllMarkers* Seurat function.

**Table S5: Differential Chromatin Accessibility**

This table contains the output from differential chromatin accessibility analysis performed using the DESeq2 R package. The columns include the **Peak ID** which describes the hg19 coordinates of the peak, and the standard output of DESeq2, including **baseMean**, **log2FoldChange**, **lfcSE** (standard error of the log2 fold change), **stat** (the Wald statistic), **p-value**, and the Benjamini Hochberg **Adjusted p-value**. Non-significant (adjusted p-value > 0.05), IFN-γ upregulated (adjusted p-value < 0.05 & log2FoldChange > 0) and IFN-γ downregulated (adjusted p-value < 0.05 & log2FoldChange < 0) peaks are reported.

**Table S6: ATAC-seq Gene Set Enrichment with GREAT**

This table contains the results from the GREAT Ontology ATAC-seq peak enrichment analysis. The columns include **Database** (Molecular Function, Cellular Component or Biological Process), the gene set or **Term** and the **GO ID**, the **FoldEnrichment, Pvalue**, and the Benjamini Hochberg Adjusted p-value (**FDR**) value.

**Table S7: Differential Transcription Factor Binding with TOBIAS**

This table contains the results from the TOBIAS differential transcription factor binding analysis. The columns describe the **UDID** (unique differentiation identifier) for the CVPC line, the **MotifID** from HOCOMOCO database, and the **FoldChange** and **Pvalue** computed by TOBIAS.

**Table S8: Chromatin State Enrichment**

This table contains the results from differential chromatin state enrichment of IFN-γ upregulated ATAC-seq peaks in 833 EpiMap tissues. The columns contain the **BSS ID** and **EpiMap Tissue Name**, the tested **Collapsed Chromatin State**, the Odds Ratio (**OR**), the **Pvalue** and Benjamini Hochberg Adjusted P-value (**Padj**) from the Fisher’s Exact test, whether or not the corresponding tissue was **Plotted** in Figure 3c. Additional information about the tissues can be obtained from the EpiMap metadata file (https://personal.broadinstitute.org/cboix/epimap/metadata/Imputation_Metadata.xlsx).

**Table S9: GWAS variant-ATAC peak Overlap**

This table contains two sheets corresponding to the GWAS overlap analysis.

Sheet 1 is a manifest with the Pan-UKB metadata for the 9 cardiac disease GWAS analyzed in this study. It contains a **Description** of the cardiac disease, the Pan-UKB **Filename**, and the **wget** command used to download the summary statistics.

Sheet 2 reports the 938 GWAS variants that overlapped a CVPC ATAC-seq peak. It contains **Trait_Description** and **Trait_ID** columns that are the Pan-UKB descriptors for the cardiac disease, the **Variant_ID** and **RSID** for the overlapping GWAS variants, the identifier for the overlapped CVPC ATAC-seq peak (**Peak_ID**), and a **Differential_Accessibility** column which describes whether the GWAS variant is in an IFNγ-upregulated (“Upregulated”) peak or a peak that is not differentially accessible (“Not DE”).

